# Real-Time and Site-Specific Perturbation of Dynamic Subcellular Compartments Using Femtosecond Pulses

**DOI:** 10.1101/2025.02.19.639204

**Authors:** Seohee Ma, Bin Dong, Matthew G. Clark, R. Michael Everly, Shivam Mahapatra, Chi Zhang

**Affiliations:** James Tarpo Jr. and Margaret Tarpo Department of Chemistry, Purdue University, 560 Oval Dr., West Lafayette, IN 47907, USA; Purdue Institute for Cancer Research, 201 S. University St., West Lafayette, IN 47907, USA; Purdue Institute of Inflammation, Immunology, and Infectious Disease, 207 S. Martin Jischke Dr., West Lafayette, IN 47907, USA

## Abstract

Understanding laser interactions with subcellular compartments is crucial for advancing optical microscopy, phototherapy, and optogenetics. While continuous-wave (CW) lasers rely on linear absorption, femtosecond (fs) lasers enable nonlinear multiphoton absorption confined to the laser focus, offering high axial precision. However, current fs laser delivery methods lack the ability to target dynamic molecular entities and automate target selection, limiting real-time perturbation of biomolecules with mobility or complex distribution. Additionally, existing technologies separate fs pulse delivery and imaging, preventing simultaneous recording of cellular responses. To overcome these challenges, we introduce fs real-time precision opto-control (fs-RPOC), which integrates a laser scanning microscope with a closed-loop feedback mechanism for automated, chemically selective subcellular perturbation. fs-RPOC achieves superior spatial precision and fast response time, enabling single- and sub-organelle microsurgery of dynamic targets and localized molecular modulation. By applying a pulse-picking method, fs-RPOC independently controls laser average and peak power at any desired subcellular compartment. Targeting mitochondria, fs-RPOC reveals site-specific molecular responses resulting from fs-laser-induced ROS formation, H_2_O_2_ diffusion, and low-density plasma generation. These findings offer new insights into fs laser interactions with subcellular compartments and demonstrate fs-RPOC’s potential for precise molecular and organelle regulation.

## Introduction

Understanding how cells and subcellular compartments respond to light interactions is essential for advancing optical microscopy, phototherapy, and optogenetics. Light-matter interaction mechanisms in biological systems vary with wavelength. Ultraviolet (UV) light can generate reactive oxygen species (ROS) or directly damage DNA^1^, while mid-infrared photons primarily cause photothermal effects by matching the vibrational absorption energy of biomolecules^2^. The near-infrared (NIR) region is particularly advantageous for biological imaging due to reduced photon absorption and minimal off-target effects. Pulsed lasers, particularly femtosecond (fs) lasers, interact with biomolecules in ways distinct from continuous-wave (CW) lasers. While biomolecules exhibit relatively weak linear absorption of NIR fs pulses, the high peak power of these laser pulses facilitates multiphoton absorption, a nonlinear process that can excite fluorescent molecules, generate ROS, induce localized heating, and activate photo-responsive proteins exclusively at the laser focus^3,4^. At higher peak power, fs laser can lead to multiphoton ionization and low density plasma (LDP) generation, resulting in more pronounced perturbations to local molecules^3,5^.

Low-energy fs lasers are widely used in biological imaging modalities, such as multiphoton-excitation fluorescence (MPEF)^6^, transient absorption^7^, and stimulated Raman scattering microscopy^8^. NIR fs pulses enable deeper tissue penetration with minimal impact to objects outside the focal plane^6^. Nonlinear optical effects further ensure that optical signals from multiphoton processes are restricted to the focal plane, providing intrinsic optical sectioning capabilities. Beyond imaging, fs pulses have been used to induce localized perturbation to biological samples. For example, fs laser pulses targeted to the endoplasmic reticulum have been shown to trigger calcium leakage into the cytosol^9-12^. Fs lasers have also been employed for nanosurgery of the cell wall, induction of cell fusion^13^, disruption of mitochondrial functions^14-17^, and cutting of actin fibers^17,18^. Additionally, fs laser pulses are used to activate optogenetic proteins^19,20^ and release small molecules into subcellular compartments. Remarkably, fs laser treatment has been demonstrated to enhance neuronal regeneration^21-23^.

Despite their extensive use in regulating and stimulating biological processes, fs lasers have yet to achieve their full potential due to limitations in delivery and dosage control. Conventional methods rely on static imaging followed by manual laser targeting, which is insufficient for dynamic molecular entities in live cells. These approaches lack automated target selection and flexible dosage control, making it impossible to simultaneously control multiple or complex molecular targets. Moreover, they separate optical control from imaging, preventing real-time monitoring of cellular responses during fs perturbations. These limitations leave critical gaps in understanding fs laser effects on biomolecules and organelles.

Here, we introduce fs real-time precision opto-control (fs-RPOC), a technology that integrates fs laser pulses with a closed-loop feedback system on a laser-scanning microscope for fully automated, site-specific, and chemically selective subcellular perturbation with fs laser pulses. Unlike conventional methods, fs-RPOC facilitates simultaneous optical perturbation and imaging for any complex or mobile molecular targets and enables real-time observation of cellular responses during fs laser perturbations. Compared to previous RPOC utilizing CW lasers^24,25^, fs-RPOC achieves unparalleled axial precision in a three-dimensional (3D) volume by leveraging multiphoton absorption. Additionally, a pulse-picking method integrated with RPOC allows independent tuning of the fs laser’s average and peak power, facilitating diverse, on-demand light-matter interaction mechanisms. Fs-RPOC enables high-precision single- and sub-organelle microsurgery and real-time perturbation of dynamic molecular entities, even in highly mobile systems. Applying fs-RPOC to precisely perturb mitochondria, we uncovered distinct site-specific cellular responses across varying laser parameters, induced by localized ROS formation, LDP generation, and delocalized H_2_O_2_ diffusion. Time-dependent and site-specific fluorescent molecule changes reveal the intricate impact of fs laser pulses and their induced chemical stimuli on intracellular molecules.

By advancing our ability to perturb biomolecules with fs laser pulses at high spatial precision and simultaneous imaging, fs-RPOC opens new avenues for understanding local or global cell responses to site-specific molecular activities.

## Results

### The femtosecond RPOC system

**Figure 1A** shows a schematic of the fs-RPOC system, which allows for the selection of four CW lasers for optical signal excitation and opto-control. In this study, only the 473 nm and 589 nm lasers were utilized to excite fluorescence signals from fluorescent proteins or organic dyes. Fluorescence detection is carried out using two photomultiplier tubes (PMTs). These fluorescence signals, originating from biomolecules or organelles, are used to identify active pixels (APXs), defined as the pixels where action lasers are activated^26^. A fs laser source (InSight X3+, Spectral Physics) serves as the major action laser, enabling the perturbation or control of biomolecular activities. The 1045 nm fs laser, with a pulse width of 150 fs, is directed to an acousto-optic modulator (AOM) for simultaneous pulse-picking and RPOC.

**Figure 1.**
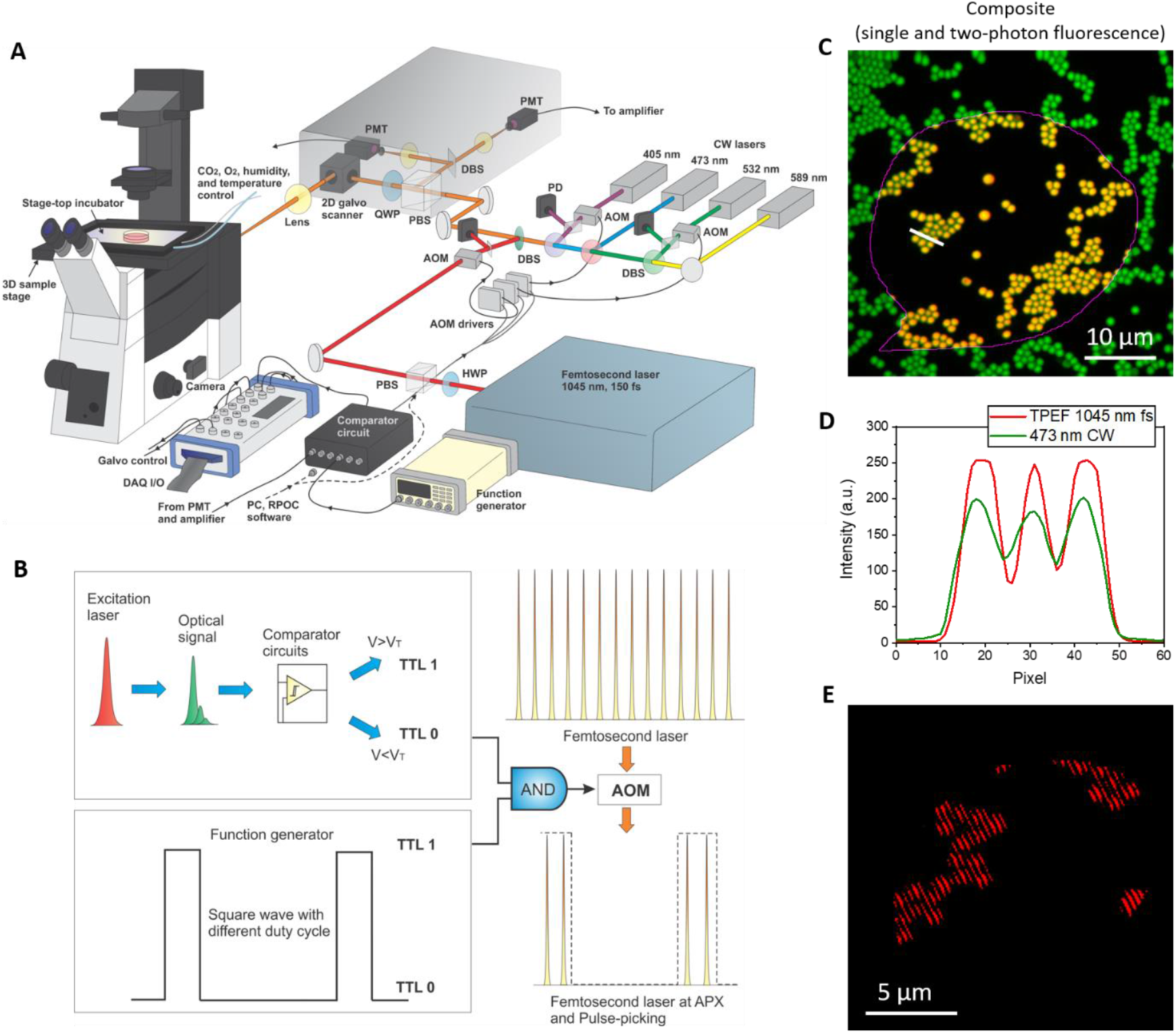
The fs-RPOC technology. (A) Schematic diagram illustrating the optical and electronic configurations of the fs-RPOC system. Abbreviations: AOM, acousto-optic modulator; PBS, polarization beam splitter; PD, photodiode; DBS, dichroic beam splitter; PMT, photomultiplier tube; DAQ, data acquisition system; HWP, half-wave plate; QWP, quarter-wave plate. (B) Conceptual illustration of achieving both RPOC and pulse picking using a single AOM. (C) Fluorescence imaging of 1 µm fluorescent polymer particles using two modalities: two-photon excitation fluorescence (TPEF) with 1045 nm laser pulses (red) and single-photon excitation fluorescence with a 473 nm continuous-wave (CW) laser (green). The RPOC software was used to select regions of interest (ROIs) for TPEF excitation with the 1045 nm laser. Yellow indicates overlapping signals from the two modalities. (D) Fluorescence intensity profiles for both modalities along the line in panel C. (E) TPEF signals from the same field of view (FOV) as in panel C, acquired under 30 kHz intensity modulation of the 1045 nm fs excitation laser at a 50% duty cycle.

**Figure 1B** illustrates the concept of using the same AOM for both RPOC and pulse-picking functionalities. The RPOC function is achieved by actively comparing optical signals from the sample to a preset threshold using a comparator circuit in real time^24,26^. Pulse picking is achieved by varying the duty cycle of a square wave sent to the AOM^27^. A digital AND operation combines the comparator circuit output with the square wave, generating transistor-transistor logic (TTL) commands for the AOM for simultaneous RPOC and pulse picking. In addition to relying solely on the comparator circuit, interactive RPOC software can be used to manually outline regions of interest (ROI) within the field of view (FOV). This software facilitates the simultaneous application of different optical treatment conditions or allows the selection of specific molecular targets for treatment^25^.

The 1045 nm fs laser operates at 80 MHz with a maximum output power of 3.5 W and a pulse width of approximately 150 fs. After passing through the optical components before the microscope, the measured pulse width at 1045 nm is ∼250 fs. A square wave with a frequency of 1 MHz and a tunable duty cycle ranging from 1.33% to 97% was used for pulse-picking. Accounting for power losses from the optical components in the system, the maximum average power and peak power that can be delivered to the sample are approximately 330 mW and 16.5 kW, respectively.

To ensure precise spatial overlap of the CW excitation lasers and the 1045 nm fs laser in both lateral and axial dimensions, the platform was tested using a sample containing 1 μm fluorescent polymer microparticles. The single-photon fluorescence signals excited by the 473 nm CW laser (green) and the two-photon excitation fluorescence signals from the fs laser (red) showed excellent spatial overlap (yellow) in both lateral and axial dimensions (**Figures 1C, D, S1**).

To validate the system’s capability for simultaneous RPOC and pulse-picking, a 30 kHz square wave with a 50% duty cycle was applied while an ROI was manually selected using the RPOC software. The square wave frequency produced a distinct intensity modulation pattern within the ROI, evidenced by patterned APXs from the fluorescent microparticles as shown in **Figure 1E**, confirming the successful integration of pulse-picking within the RPOC framework. This implementation enables independent control of the laser’s average and peak power, facilitating the evaluation of their respective impacts on cellular responses.

### Femtosecond RPOC confines laser perturbation to the focal plane for single- and sub-organelle microsurgery

Although optical control or treatment using RPOC with CW lasers provides high spatiotemporal precision^24^, the perturbation mechanism is based on linear light absorption, which can affect cells or organelles outside the focal plane. In contrast, fs laser-based optical perturbation relies on nonlinear optical effects, which are highly localized to the laser focus where the energy density is highest (as illustrated in **Figures 2A and 2B**). This results in enhanced 3D precision, particularly along the axial direction, while minimizing the impact on out-of-focus planes. To compare the effects of optical treatment using a 405 nm CW laser and a 1045 nm fs laser, we performed 3D RPOC and simultaneously fluorescence imaging of cells cultured in 3D structures. HeLa cells transfected with EB3-EGFP were used to visualize cellular EB3 proteins. A focal plane was selected for laser treatment, and the impacts of RPOC on this plane and a different axial layer were evaluated (**Figure 2C**). The RPOC software was applied to select APXs within a cell at the interaction plane^25^.

**Figure 2.**
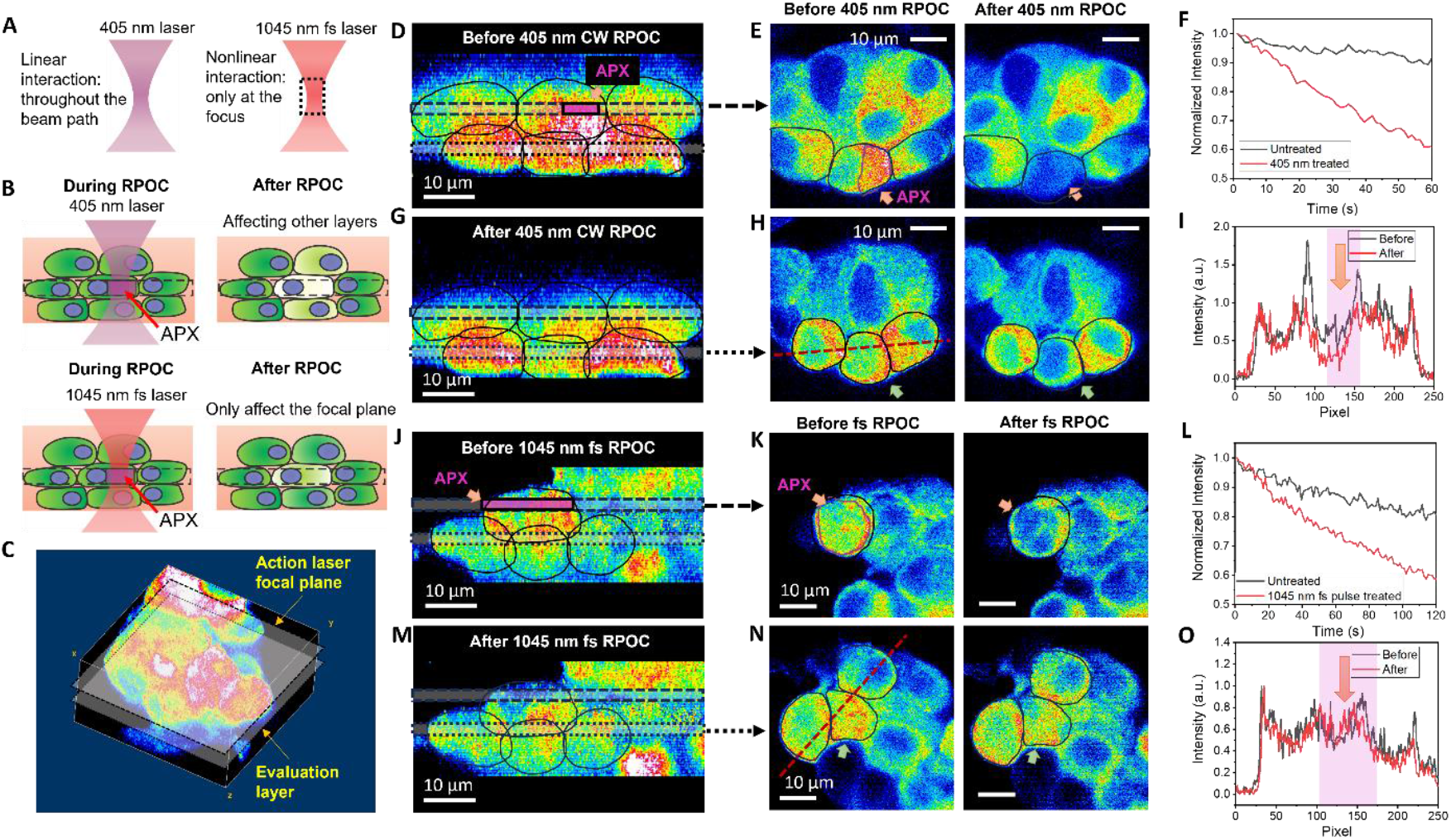
Enhanced axial precision with fs-RPOC. (A) Illustration of linear interactions occurring throughout the beam path with continuous-wave (CW) lasers versus nonlinear interactions confined to the laser focus with fs lasers. (B) Schematic comparison showing extended off-focus impact on cells using CW lasers versus minimal off-focus impact with fs near-infrared (NIR) lasers. (C) Example of a 3D fluorescence image depicting cells, the action laser focal plane, and an evaluation plane located below the interaction focal plane. (D) Side view of the 3D live HeLa cell image showing EB3-EGFP fluorescence signals prior to treatment with a 405 nm CW laser. Cells are outlined in the images. Dashed and dotted areas indicate the action laser focal plane and the evaluation plane, respectively. (E) Top view of the interaction focal plane before and after RPOC. APXs selected by the RPOC software are outlined in the image. (F) Fluorescence signal decay comparison between treated and untreated cells. (G) Side view of the 3D image showing EB3-EGFP signals after 405 nm CW laser treatment. (H) Top view of the evaluation plane before and after RPOC. (I) EB3-EGFP fluorescence signals from a cell (green arrow in panel H) below the treated cell before and after treatment with a 405 nm laser at 440 µW. (J–O) Similar to panels D–I, but with the selected cell treated using 13 mW fs laser pulses at 1045 nm at APXs. A negligible signal decrease was observed in the cell beneath the treated cell.

When the 405 nm CW laser was applied (0.44 mW on the sample, 0.25 mJ total dose received by the cell across 68 s), the treated cell exhibited a significant loss of EB3 signal compared to untreated controls (**Figures 2D–2F**). Furthermore, the 405 nm laser affected the cell beneath the treated cell (**Figures 2G–2I**). This cell, located below the treatment laser focal plane, showed a pronounced decay in EB3-EGFP signals after treatment (**Figure 2I**). In comparison, treatment with a 1045 nm fs laser (13 mW, 80 MHz, 650 W peak power, 25.8 mJ total dose received by the cell over 120 s) resulted in similar EB3-EGFP signal decay in the treated cell within the focal plane, likely due to the reduced perturbation from fs laser pulses (**Figures 2J–2L**). Notably, the cell beneath the treated cell exhibited negligible changes in EB3-EGFP signals after treatment (**Figures 2M–2O**). These findings indicate that, compared to the 405 nm CW laser, which perturbs cells via linear absorption, the 1045 nm fs laser minimizes off-focus perturbations, leading to reduced impact on cells outside the focal plane.

The high 3D spatial accuracy of fs-RPOC enables single- and sub-organelle microsurgery within a 3D volume. To demonstrate this capability, we labeled cells with MitoTracker Red and performed 3D RPOC on selected mitochondria. The results showcase both lateral and axial precision in perturbing targeted and mobile mitochondria, as illustrated in **Figure 3A**. Using the RPOC software, we simultaneously selected five ROIs and applied a 22 mW (1.1 kW peak power) fs laser at 1045 nm, in conjunction with the comparator circuit box, to treat mitochondria within the designated ROIs. The APXs within the ROIs were automatically identified based on MitoTracker fluorescence signals and a predetermined intensity threshold.

**Figure 3.**
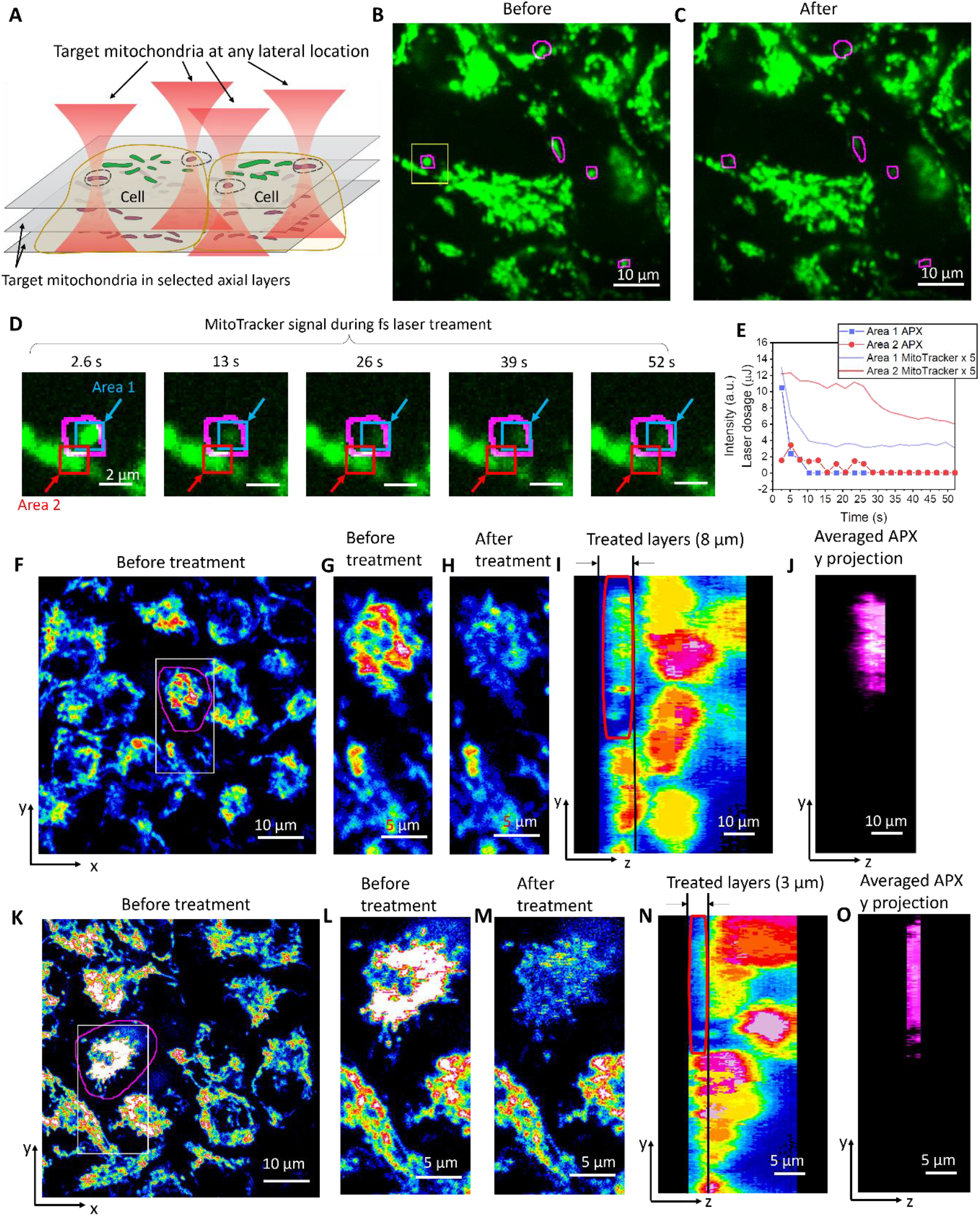
Microsurgery of mitochondria with high lateral and axial accuracy within a 3D system. (A) Illustration of precise mitochondrial perturbation with single-or sub-mitochondrion accuracy in both lateral and axial dimensions. (B) MitoTracker signals from live HeLa cells labeled with MitoTracker. Regions of interest (ROIs) selected for fs RPOC are outlined. (C) MitoTracker signals after RPOC treatment with 22 mW, 250 fs laser pulses with 1.1 kW peak power. Mitochondria within or entering the selected ROIs show significantly reduced MitoTracker signals at different time points. (D) A selected area in panel B at different times during RPOC. Area 1 indicates a fully treated mitochondrion, while Area 2 indicates a partially treated mitochondrion. (E) MitoTracker signals and APX (laser dosage) at Areas 1 and 2 in panel D. (F) MitoTracker signals from HeLa cells at one interaction layer before treatment. The ROI selected by RPOC is outlined in magenta. At any axial layer for treatment, APXs within this ROI are automatically selected based on MitoTracker signals. (G) Enlarged view of a selected area from panel D. (H) The same area as in panel E after RPOC treatment with 22 mW, 250 fs laser pulses. The treatment was performed across the bottom 8 µm of the ROI, with steps of 1 µm per frame over 8×5 frames (100 seconds). (I) Side view of MitoTracker signals after treatment. The treated ROI is outlined in red. (J) Side view of APXs automatically selected by fs RPOC within the outlined ROI. (I–M) Similar to panels D–H, but with a 3 µm layer at the bottom of the cell selected in panel I treated for 3×6 frames (48 seconds). MitoTracker signals from mitochondria within the selected volume are significantly diminished.

As shown in **Figures 3B and 3C**, fs-RPOC selectively and precisely perturbed the targeted mitochondria within the lateral plane, as evidenced by a significant reduction in MitoTracker signals following treatment. Despite the highly dynamic nature of mitochondria, RPOC enables precise targeting through automated selection of ROIs. The RPOC-selected mask and averaged APXs of this treatment are shown in **Figure S2**. Real-time APXs during treatment can be found in **Video S1**. Moreover, mitochondria that partially migrated into the ROIs during treatment were also affected, exhibiting noticeable MitoTracker signal loss (**Figure 3D, area 2**). The laser doses administered to the fully and partially treated mitochondria in **Figure 3D** were 15.0 μJ and 14.4 μJ, respectively. Notably, the fully treated mitochondria received a substantially higher laser dose at the outset (**Figure 3E**), resulting in a more rapid decline in MitoTracker signals. This signal decay is primarily attributed to ROS generated at the APXs, which oxidize MitoTracker, as well as the leakage of MitoTracker into the cytosol due to mitochondrial membrane damage. In contrast, the partially treated mitochondria were exposed to a lower fs laser dose over an extended period, leading to a slower initial decline in MitoTracker signals (**Figure 3E**). However, this sequential exposure eventually triggered a more rapid signal decrease after reaching a threshold at approximately 25 seconds, despite the termination of APXs and optical treatment at this time point. The gradual signal reduction is largely driven by hydrogen peroxide (H_2_O_2_) diffusing from the APX into other parts of the mitochondrion, while the accelerated decline around 25 seconds is likely due to mitochondrial membrane damage, causing MitoTracker to leak into the cytosol. Other ROS species, in comparison, have much shorter lifetimes and limited diffusion ranges compared to H_2_O_2_. These findings suggest that fs laser treatment targeting a small region of the mitochondrion can ultimately compromise the integrity and function of the entire organelle.

Furthermore, we applied RPOC to select APXs within a 3D cell structure attached to a culture dish (8 µm thickness) in a 3D population. After 104 seconds of laser scanning, mitochondria within the targeted cell exhibited a significant reduction in MitoTracker signals (**Figures 3F-J**), while mitochondria in neighboring cells remained unaffected. The APXs during treatment at each axial plane and time are shown in **Figure S3**, which are automatically selected by RPOC based on MitoTracker signals at each layer and time point. By narrowing the treatment layer to a 3 µm thickness, as shown in **Figures 3K-O**, we achieved selective perturbation of mitochondria within the specified layers of the targeted cell, without disturbing other organelles in the treated area and mitochondria in other layers of the same cell.

The observed decrease in MitoTracker signals is likely attributed to a combination of two-photon photobleaching, local ROS-induced oxidation, and dye leakage caused by mitochondrial membrane damage. LDP formation and laser breakdown processes are unlikely to occur at the applied low peak power range. These results underscore the capability of fs RPOC to precisely target cellular organelles or other entities with high 3D spatial precision, avoiding unintended effects on neighboring locations in both lateral and axial directions.

### Femtosecond laser interaction with cells

Interactions of pulsed lasers with solid or liquid samples have been explored both theoretically and experimentally. To perform effective cellular perturbation or ablation, two schemes are typically applied. The first is using low repetition rate picosecond or nanosecond lasers, which have µJ to mJ pulse energy and perturb samples majorly through heating-induced laser breakdown, shock wave, or intense plasma generation^28-31^. The second scheme is by applying lower energy laser pulses but with short pulse duration in the fs range at a relatively high repetition rate^15,23,32-34^. Mechanisms involved in such fs laser interactions can range from multiphoton-mediated electron transfer to LDP generation and ionization below the laser breakdown threshold^3^. Notably, the tuning range of fs laser interaction with samples is broad, spanning from minimal perturbing power used in nonlinear optical imaging to extensive ionization and free electron generation reaching the laser breakdown level^3^.

From previous theoretical and experimental studies, the radiation power on the level of 10^11^-10^12^ W/cm^2^ in the NIR range is found to be on the margin of generating detectable perturbations to live cells. This power range was found to generate 1 free electron per pulse or perturb functions of mitochondria^3^.

To compare our work with previous studies from other groups, we perform a calculation of the radiation in our RPOC system. Assuming the fs laser has a 33 mW average power on the sample with a 250 fs pulse duration, the pulse energy and peak power on the sample at 80 MHz are

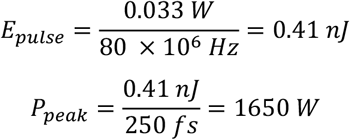

In this work, we are using an objective lens with a 1.2 numerical aperture (NA). The focal spot size at the diffraction limit gives a diameter of 531 nm. Therefore, the laser focus has an area of about 0.22 µm^2^. The radiation power at the focus is calculated to be

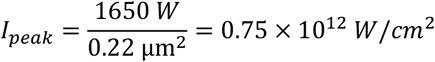

This range is far below the laser breakdown radiation level^3^, but on the margin of perturbing cell functions^3^. To increase peak power while minimizing heat generation, our pulse-picking method could separately tune the average and peak power of fs laser pulses^27^. For example, when increasing the laser power to 330 mW and applying a 10% duty cycle in pulse peaking, we could achieve a 33 mW average power on the sample with a peak power of 16.5 kW. Such a peak power is close to 10^13^ W/cm^2^, which was reported to induce fast mitochondrion ablation^3,35^. At this peak power, LDP can also potentially be generated, mediated by intrinsic biomolecules such as nicotinamide adenine dinucleotide hydrogen (NADH), flavin adenine dinucleotide (FAD), and cytochrome c, or by exogenous fluorophores introduced into the cellular system. In this work, by using the pulse-picking method, we can achieve various combinations of laser average and peak power as summarized in Table 1.

**Table 1.** The laser average power, peak power, and duty cycle that were applied in this study.

| Average power, no pulse picking, before microscope | Duty cycle for pulse picking | Average power on the sample (22% transmission rate) | Peak power on the sample |
| --- | --- | --- | --- |
| 1.5 W | 10% | 33 mW | 16.5 kW |
| 1.5 W | 5% | 16.5 mW | 16.5 kW |
| 1.5 W | 1.33% | 4.4 mW | 16.5 kW |
| 0.5 W | 30% | 33 mW | 5.5 kW |
| 0.5 W | 15% | 16.5 mW | 5.5 kW |
| 0.5 W | 4% | 4.4 mW | 5.5 kW |
| 0.1 W | 75% | 16.5 mW | 1.1 kW |
| 0.1 W | 20% | 4.4 mW | 1.1 kW |

### The dependence of average and peak power of laser pulses

At low average power, NIR fs lasers interact with subcellular organelles mostly through multiphoton processes. Using fs-RPOC, we investigated the effects of varying laser power levels on cellular responses. A schematic representation of fs laser interaction with MitoTracker-labeled mitochondria at APXs is shown in **Figure 4A**. During laser interaction at APXs, multiphoton absorption, primarily mediated by MitoTracker or/and inherent biomolecules, may result in photobleaching of MitoTracker by directly altering the structure of the dye. Additionally, this absorption generates ROS, which induces local oxidation of MitoTracker, further reducing its fluorescence signals. Under very high peak power conditions, multiphoton absorption can also lead to the formation of LDPs, which both directly degrade MitoTracker molecules or indirectly contribute and their damage via LDP-induced ROS.

**Figure 4.**
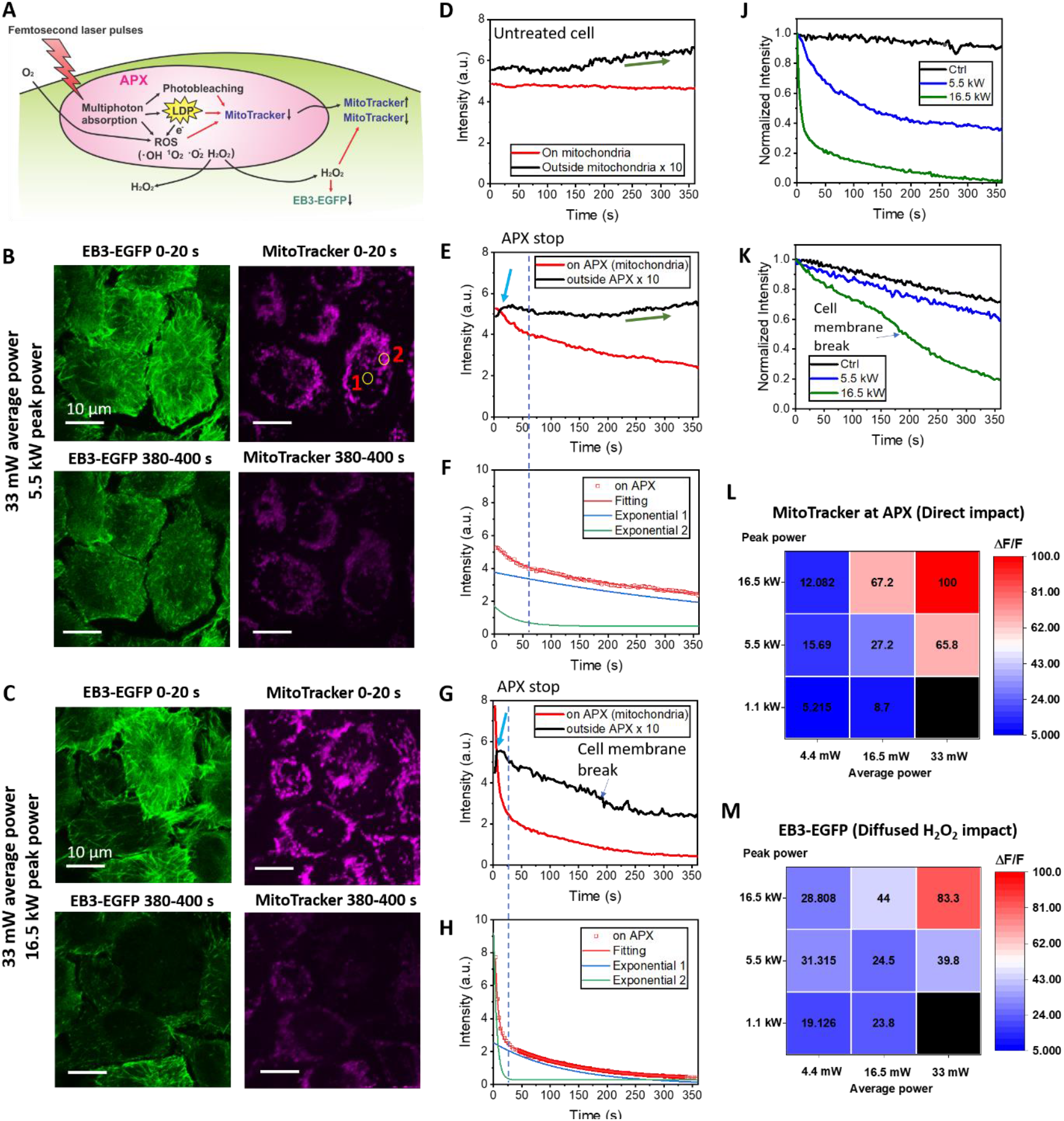
Cellular responses to fs laser treatment of mitochondria at varying average and peak power levels. (A) Schematic illustration of the mechanism by which fs lasers interact with molecules within and outside APXs. (B) EB3-EGFP and MitoTracker signal at the start and end of RPOC. APXs are identified using MitoTracker signals localized to mitochondria. Regions of interest (ROIs) outside APXs (1) and within APXs (2) are selected for time-lapse signal analysis. The fs laser pulses have 33 mW average power and 5.5 kW peak power. (C) Similar to panel B but using 33 mW average power and 16.5 kW peak power fs laser pulses. (D) Time-lapse MitoTracker signals on and outside mitochondria in untreated cells. (E) Time-lapse MitoTracker signals on and outside APXs in HeLa cells from panel B. (F) Dual-exponential fitting of time-lapse MitoTracker signals within APXs in panel E. The blue arrow indicates the initial rise of MitoTracker signals outside APXs. The dotted line marks the automatic termination time of APXs due to declining MitoTracker signals. (G, H) Similar to panels E and F, but for cells treated with 16.5 kW peak power, as shown in panel C. (J) Average MitoTracker signals from untreated and treated HeLa cells in panels B and C, with peak powers of 5.5 kW and 16.5 kW, respectively. (K) Similar to panel J, but for EB3-EGFP signals under the same conditions. (L) Decay rate (percentage) of MitoTracker signals after RPOC using different average and peak power values. (M) Decay rate (percentage) of EB3-EGFP signals after RPOC using fs laser pulses with different average and peak power.

Damage to mitochondrial membranes caused by these mechanisms can lead to the leakage of MitoTracker from mitochondria, increasing MitoTracker signals in the vicinity of the APXs. Additionally, H_2_O_2_, a byproduct of ROS generated at the APXs, can diffuse from the mitochondria into the cytosol^36^. In contrast, other ROS species such as singlet oxygen, hydroxyl radicals, and superoxide ions are less likely to diffuse significantly beyond the APXs^37,38^. The diffused H_2_O_2_ can oxidize both EB3-EGFP and the leaked MitoTracker outside the APXs^39^. The competing effect of MitoTracker leakage from APXs and its oxidation by H_2_O_2_ could lead to varying changes in MitoTracker signal outside APXs over time.

The efficiency of multiphoton processes depends on peak power levels, leading to significant variations in fs laser-induced interactions at APXs and the resulting cellular responses. To explore these differences, we employed fs-RPOC with pulse-picking to independently control laser average and peak power, targeting mitochondria in EB3-EGFP-transfected HeLa cells. APXs were automatically selected based on MitoTracker signals using comparator circuits of the RPOC system. The use of comparator circuits enables real-time dynamic comparison of MitoTracker signals against a predefined threshold. As the MitoTracker signals decay during treatment, the number of APXs and treatment laser dose typically decreases over time. The treatment automatically terminates once the MitoTracker signals fall below the threshold. Changes in EB3-EGFP signals primarily reflect the impact of H_2_O_2_ released from APXs, while MitoTracker signals primarily indicate both direct laser interactions and indirect effects of H_2_O_2_ (**Figures 4B-K**). We monitor both effects using separate fluorescent channels.

Using pulse-picking, we first achieved an average power of 33 mW for 1045 nm laser pulses, with a peak power of 5.5 kW at APXs. Over a 380-second RPOC period, EB3 comets exhibited significant shortening (**Figure 4B, Video S2**), indicating strong phototoxicity induced by fs laser pulses at APXs^40^. When the peak power was increased to 16.5 kW while maintaining the same 33 mW average power, the fs laser treatment caused more severe photodamage, characterized by membrane rupture and the complete loss of intracellular EB3-EGFP and MitoTracker signals (**Figure 4C, Video S3**).

To quantify these effects, we first monitored MitoTracker signals on and outside mitochondria in untreated cells as a control. On mitochondria, MitoTracker signals remained stable over 380 s of imaging (**Figure 4D**). Outside mitochondria, signals showed a slow increase, likely due to MitoTracker leakage induced by the 473 nm imaging laser (**Figure 4D**).

We selected areas (for example, **Figure 4B**, areas 1 and 2) at APXs and outside APXs to understand the direct fs laser impact and the indirect MitoTracker leaking and oxidation by H_2_O_2_. In cells treated with 33 mW average power and 5.5 kW peak power, MitoTracker signals at APXs show a decay (**Figure 4E**), highlighting the direct effects of fs laser pulses. This decay followed a two-phase pattern (**Figure 4F**): an initial rapid decline due to APX activity, ceasing around 20 seconds as MitoTracker signals fell below the APX threshold, followed by a slower decay caused by H_2_O_2_oxidation and MitoTracker leakage from mitochondria. Outside APXs, MitoTracker signals initially rose rapidly, indicating mitochondrial leakage into the cytosol, then declined due to H_2_O_2_diffusion and oxidation of leaked MitoTracker (**Figure 4E**). Once the fs laser at APXs was automatically deactivated, the production of H_2_O_2_ stopped, leading to partial recovery of MitoTracker signals outside APXs due to the continued dye leakage, similar to the untreated cells (**Figure 4D**).

Increasing the fs laser peak power to 16.5 kW (while keeping the same 33 mW average power) significantly amplified these effects. In the MitoTracker channel, APX signals decayed rapidly during laser activation, followed by a slower decay post-deactivation (**Figure 4G**), both well fitted by two-phase exponential functions (**Figure 4H**). This decay occurred more quickly than in the 5.5 kW peak power case, with MitoTracker signals dropping below 15%, indicating extensive molecular damage at APXs. Outside APXs, signals initially spiked before declining more rapidly than in the 5.5 kW case due to higher H_2_O_2_ production and leakage. Unlike the 5.5 kW case, which exhibited a partial increase in the MitoTracker signal, the 16.5 kW peak power case showed continued signal decay outside APXs even after APX deactivation, suggesting elevated cytosolic H_2_O_2_ levels. Higher H_2_O_2_ generation at 16.5 kW peak power led to significantly lower MitoTracker levels in mitochondria, preventing further MitoTracker signal rise due to leakage. Furthermore, around 170 s, plasma membrane rupture further accelerated MitoTracker leakage outside of the cell (**Figure 4G**). The averaged MitoTracker signals change over time for both treatment conditions, along with the control, are compared in **Figure 4J**.

In the EB3-EGFP channel, treatment with 5.5 kW peak power resulted in less signal decay compared to 16.5 kW (**Figure 4K**), confirming greater H_2_O_2_ generation at higher peak power. Additionally, cell membrane rupture was also detectable as the accelerated EB3 signal loss at around 170 s, consistent with the MitoTracker response in **Figure 4G**.

The distinct fluorescent signal changes observed at different laser peak power levels demonstrate that fs laser-induced perturbations to mitochondria and cellular functions are highly peak power-dependent. Varying peak power levels result in differences in the rates of molecular damage at APXs, leakage from APXs, and oxidation outside APXs. These effects are reflected in the fluorescence signal dynamics over time.

To further assess the impact of fs laser average and peak power on mitochondrial-targeted cellular responses, we compared the fluorescence signal decay rates (ΔF/F) in the EB3-EGFP and MitoTracker channels across treated cells over 380 s using different combinations of average and peak power (**Figures 4L and 4M**). At 5.5 kW peak power, significant effects emerged when the average power reached 33 mW, whereas at 16.5 kW peak power, notable impacts were observed at average powers above 16.5 mW. These results indicate that at 16.5 mW average power and above, the cellular response is peak power dependent. Consequently, the decline in cellular fluorescence signals under high peak power fs laser pulses is unlikely to result from thermal effects due to linear absorption.

This conclusion is further supported by experiments illuminating the entire FOVs with similar average power but much lower peak power, as shown in **Figure S4**. Compared to the control group (no fs laser applied), illuminating the entire FOV over 320 seconds with laser average powers of 11 mW or 22 mW and peak powers ranging from 344 to 688 W resulted in minimal EB3-EGFP signal decay. Higher peak power only slightly enhances the shortening of EB3 comets (**Video S4**). When centrosomes were present within the FOV and illuminated by fs lasers, the corresponding cells showed slightly enhanced shrinkage. These findings further confirm that at high peak power, fs laser-induced perturbations to cellular functions are primarily due to multiphoton processes rather than linear absorption of NIR fs laser pulses.

The results in **Figure 4** indicate that cellular responses depend on both peak and average power. While high peak power is essential to enable efficient multiphoton absorption, the generated ROS or LDP must surpass a critical threshold to elicit significant cellular effects. At lower concentrations, ROS or LDP produced through multiphoton processes at APXs do not induce noticeable cellular responses within the observed timeframe. Future studies will explore long-term cellular responses under varying laser power levels and dosages.

### The involvement of LDP

In **Figure 4A**, LDP may be induced by multiphoton absorption of fs lasers at high peak power. These LDP are highly reactive and likely interact with surrounding molecules instantaneously. White light generation from LDP is observable only when their concentration reaches a sufficient threshold to the avalanche ionization level^3^. When using fs-RPOC to target and perturb mitochondria, we did not observe white light generation due to the MitoTracker signal decay which rapidly turned off APXs and fs laser pulses.

To confirm the involvement of LDP, we turned the action laser on throughout the entire frame using fs laser pulses with 16.5 mW average power and 16.5 kW peak power and treated the cells for 320 s. This resulted in significantly more pronounced cell damage (**Figure 5A**) compared to previous cases. Membrane rupture of Cell 1 occurred at approximately 64 seconds. Initially, EB3 comets remained intact, followed by a rapid decline in fluorescence signals and eventual plasma membrane rupture (**Figure 5A**). This cell death process differs from the mitochondria-targeted case shown in **Figure 4B**, showing no shortening of EB3 comets but rapid bleaching of EB3-EGFP, followed by membrane rupture and leakage of cellular contents. (**Video S5**).

**Figure 5.**
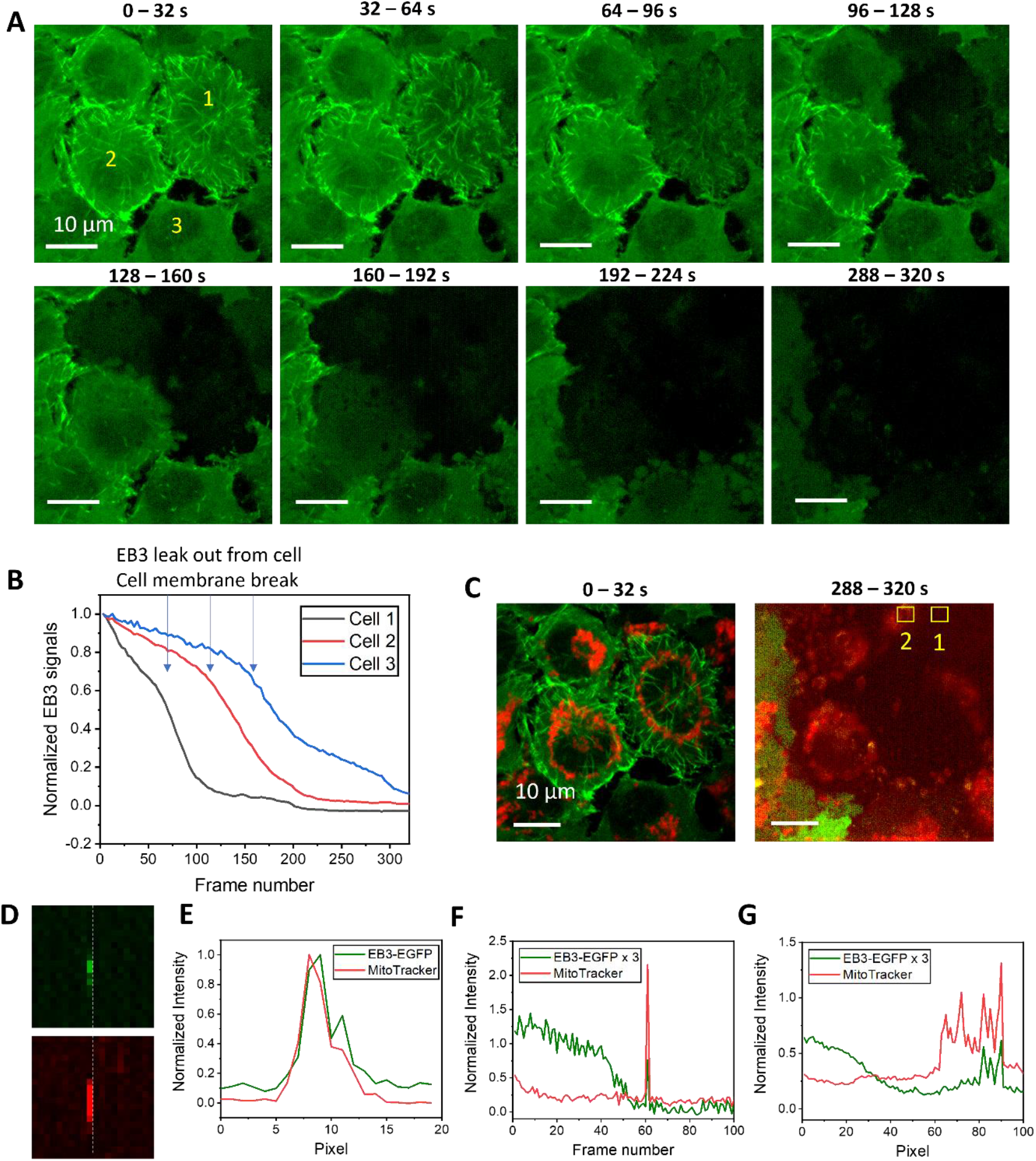
Confirming low-density plasma (LDP) involvement through white light generation. (A) EB3-EGFP signals from HeLa cells during RPOC treatment with 16.5 mW average power and 16.5 kW peak power. The fs laser is activated across the entire field of view (FOV). (B) EB3-EGFP signal decay over treatment time for three selected cells in panel A. Arrows indicate time points where plasma membrane rupture occurs. (C) Correlated EB3-EGFP and MitoTracker images from the same FOV as panel A, captured at the beginning and end of RPOC. Areas of white light generation are outlined in yellow squares. (D) White light emission detected in both EGFP and MitoTracker channels from area 1 in panel C at 61 s. (E) Intensity profiles of the white light signals in panel D. (F) Time-lapse intensity within selected area 1 from panel C in both fluorescence channels. (G) Time-lapse intensity within selected area 2 from panel C.

Additionally, due to the slight laser energy inhomogeneity within the FOV, cells exhibited photo-damage at slightly different rates. All three selected cells in the image showed membrane rupture at different time points, characterized by accelerated EB3-EGFP signal loss (**Figure 5B**) similar to observations in **Figure 4K**. During these damage processes, white light generation was observed (**Figures 5D-G**), with significantly stronger signals than fluorescence, detectable in both the EGFP and MitoTracker channels. These emissions show sharp spikes, originating from LDP-induced white light generation, and have strong spatial and temporal correlation across both fluorescence channels.

These findings indicate that 1045 nm fs laser pulses at 16.5 kW peak power can induce LDP. The rate of LDP generation and its impact on cells are dosage-dependent. At low LDP dosages, cellular functions remain largely unaffected, as shown in **Figure 4**.

## Discussion

In this study, we utilized 1045 nm fs laser pulses for cellular organelle perturbation. Our system also includes a frequency-tunable laser, which can be employed as an action laser for RPOC. This capability would enable wavelength-dependent evaluations of site-specific perturbations to cellular functions and provide insights into laser interaction mechanisms by comparing wavelength-dependent responses with the absorption spectra of specific molecules. Future studies will explore these wavelength-dependent effects.

Fs RPOC offers optimal spatiotemporal precision, particularly in the axial direction, to perturb biomolecular functions accurately. The integration of a pulse-picking mechanism into RPOC enables independent control of pulse average and peak power at APXs, allowing precise modulation of cellular targets with separately adjustable peak power levels. We demonstrated the precise targeting of selected mitochondria in a 3D system and evaluated the effects of varying average and peak power on cellular responses during mitochondrial perturbation. Different power levels elicited diverse cellular responses due to variations in the production of ROS, LDP, and molecular diffusion within cells.

This study presents an advanced system for controlling and perturbing cellular targets with fs lasers, offering flexible combinations of average and peak power. We demonstrate single- and sub-organelle microsurgery in live cells with high spatial precision and simultaneous monitoring of cell responses. Additionally, our findings provide new insights into how site-specific fs laser perturbations affect cellular responses through various multiphoton-induced processes at different power levels.

## Methods

### The fs-RPOC technology

The fs-RPOC system is based on a custom-designed confocal fluorescence microscope. It incorporates four CW lasers (405 nm, 473 nm, 532 nm, and 589 nm) for fluorescence excitation and optical control. The 405 nm and 532 nm lasers are used as optional CW action lasers. Acousto-optic modulators (AOMs, P80L-0.5 with 532B-2 driver, Isomet) are installed in the 405 nm and 532 nm laser paths, functioning as fast optical shutters. These lasers are coupled to the sample via the first-order diffraction. The beams are combined collinearly using dichroic beam splitters (DMLP567, DMLP505, DMLP425, Thorlabs), with neutral density filters in each path for laser power attenuation.

The 1045 nm fs laser (InSight X3+, Spectral Physics) passes through another AOM for simultaneous RPOC and pulse picking. All lasers are directed to the microscope via a 2D galvo scanner (Saturn-5 system, ScannerMAX), expanded to 9 mm by a lens pair, and focused through a water-immersed objective lens (UPLSAPO-S, 60X, NA 1.20, Olympus) mounted on an inverted microscope (Olympus IX73). TPEF images in **Figure 1** are acquired using the fs action laser by keeping the laser constantly on using the comparator circuit box.

Fluorescence signals are separated using a polarization beam splitter and a quarter-wave plate, then detected by two photomultiplier tubes (PMTs, H7422-40, Hamamatsu) through confocal pinholes (P300HK, Thorlabs). Filters enable the detection of EGFP and organelle dyes in separate channels. Preamplifiers (PMT4V3, Advanced Research Instruments Corporation) convert and amplify signals for data acquisition (PCIe-6363, National Instruments). A comparator circuit, previously reported^24^, automates APX selection.

Pulse picking is achieved with RPOC by real-time computation of RPOC TTL with a square wave from a function generator (DG1022Z, Rigol). The frequency and duty cycle of the square wave is tunable. The comparator circuit applies a digital AND operation to combine (multiply) the RPOC output with the square wave.

The RPOC software, developed in LabVIEW, integrates laser control and imaging^25^. It allows manual ROI delineation, automatic target selection based on optical signals, and precise targeting of dynamic molecular entities within the ROI. The software supports 3D RPOC and imaging across the ROI or the full FOV. APXs are monitored using photodiodes (PDA10A2, Thorlabs) which detect small fractions of the action lasers deflected by glass slides.

A stage-top incubator (WSKMX with STX-CO2O2, Tokai Hit) maintains precise humidity, temperature, and CO_2_levels for cultured cells, enabling continuous monitoring of the same FOV for over 48 hours without changes of culture medium.

### RPOC, imaging, and data processing

For cells in **Figures 2D-H**, a 3D acquisition with a thickness of 30 µm was performed prior to the RPOC treatment. An ROI within a cell in the interaction layer was selected using the RPOC software. The treatment was applied by scanning the 405 nm laser (440 μW on the sample) for 68 seconds consecutively at this layer within APXs, with simultaneous image acquisition during the process. The outlined areas of APX received 0.25 mJ total laser dosage. After the treatment, another 3D acquisition was conducted to compare pre- and post-treatment signals. Time-dependent intensity profiles were generated by integrating EB3 fluorescence signals from each frame during treatment and plotted using Origin 2021. The cells in **Figures 2J-N** were imaged under similar conditions as those in **Figures 2D-H**. The fs laser treatment was conducted using a 1045 nm fs laser with a pulse width of 250 fs for 120 s. The APX area received a total fs laser dose of 25.8 mJ.

Intensity profiles from cells in **Figure 2** were compared along the same line at the same layers before and after treatment. All 3D images were processed in ImageJ and displayed using a Jet color scheme.

In **Figures 3B and 3C**, separate ROIs were selected using the RPOC software. Comparator circuits were used in combination with the RPOC software to control dynamic mitochondria within the selected area. MitoTracker signals were used to identify real-time APXs through the comparator circuit. Treatments were applied by scanning the laser for 52 seconds, and 3D images were acquired following the same procedure as in **Figure 2**. In **Figures 3F-O**, RPOC treatment was continuously executed within the selected layer of the 3D cell structure for different time lengths indicated in the figure caption. Images were processed in ImageJ to display x-y and y-z projections, and 3D APXs were visualized in the y-z plane.

For **Figures 4 and 5**, intensity plots were generated for APXs and regions outside APXs using Origin 2021. Curve fitting with single or dual exponential functions was also performed in Origin 2021. Heatmap plots showing the percentage of fluorescence signal decay in each channel after treatment were generated and visualized using Origin 2021.

### Cell preparation and labeling

HeLa Kyoto EB3-EGFP cells were purchased from Biohippo and cultured in Dulbecco’s Modified Eagle Medium (DMEM, ATCC) supplemented with 10% fetal bovine serum (FBS, ATCC) and 1% penicillin/streptomycin (Thermo Fisher Scientific). The cells were seeded into glass-bottom dishes (MatTek Life Sciences) containing 2 mL of culture medium and incubated in a CO_2_ incubator at 37°C with 5% CO_2_. Cells were grown overnight until they reached approximately 50–70% confluence for live-cell imaging and RPOC.

For labeling with MitoTracker, the cells were incubated with MitoTracker Red CMXRos at a final concentration of 200 nM, according to the manufacturer’s instructions. After a 30-minute incubation at 37°C and 5% CO_2_, the cells were gently rinsed twice with a warm culture medium before imaging and RPOC.

## Acknowledgments

This work is supported by NIH R35GM147092.

## Author Contributions

Se.M, B.D. C.Z. and M.C. performed the experiments. Se.M. prepared biological samples. Se.M. and C.Z. analyzed the results. R.M.E designed the RPOC software. B.D. tested the software and maintained the RPOC system. C.Z. and Se.M. wrote the paper. C.Z. designed the project and obtained the funding.

## References

1 Zhang, X., Rosenstein, B. S., Wang, Y., Lebwohl, M. & Wei, H. Identification of possible reactive oxygen species involved in ultraviolet radiation-induced oxidative DNA damage. Free Radic. Biol. Med. 23, 980–985 (1997).

2 Jean, B. & Bende, T. Mid-IR laser applications in medicine. Solid-State Mid-Infrared Laser Sources, 530–565 (2003).

3 Vogel, A., Noack, J., Hüttman, G. & Paltauf, G. Mechanisms of femtosecond laser nanosurgery of cells and tissues. Appl. Phys. B 81, 1015–1047 (2005).

4 Quinto-Su, P. A. & Venugopalan, V. Mechanisms of laser cellular microsurgery. Methods Cell Biol. 82, 111–151 (2007).

5 Vogel, A., Noack, J., Huettmann, G. & Paltauf, G. in Multiphoton Microscopy in the Biomedical Sciences II. 202–216 (SPIE).

6 So, P. T., Dong, C. Y., Masters, B. R. & Berland, K. M. Two-photon excitation fluorescence microscopy. Annu. Rev. Biomed. Eng. 2, 399–429 (2000).

7 Davydova, D. y., de la Cadena, A., Akimov, D. & Dietzek, B. Transient absorption microscopy: advances in chemical imaging of photoinduced dynamics. Laser Photonics Rev. 10, 62–81 (2016).

8 Zhang, C., Zhang, D. & Cheng, J.-X. Coherent Raman scattering microscopy in biology and medicine. Annu. Rev. Biomed. Eng. 17, 415–445 (2015).

9 Tian, X. & He, H. Activation of Mitochondrial Ca2+ Oscillation and Mitophagy Induction by Femtosecond Laser Photostimulation. Bio Protoc. 12, e4369–e4369 (2022).

10 Shi, F., Wang, S., Zhu, Y. & He, H. Excitation of mitochondria-Endoplasmic reticulum Ca2+ coupling by femtosecond-Laser photostimulation. IEEE Photonics J. 12, 1–8 (2020).

11 He, H. et al. Manipulation of cellular light from green fluorescent protein by a femtosecond laser. Nat. Photonics 6, 651–656 (2012).

12 Wang, S. et al. Photoactivation of Extracellular-Signal-Regulated Kinase Signaling in Target Cells by Femtosecond Laser. Laser Photonics Rev. 12, 1700137 (2018).

13 Kohli, V., Elezzabi, A. Y. & Acker, J. P. Cell nanosurgery using ultrashort (femtosecond) laser pulses: applications to membrane surgery and cell isolation. Lasers Surg. Med. 37, 227–230 (2005).

14 Watanabe, W. et al. Femtosecond laser disruption of mitochondria in living cells. Med. Laser Appl. 20, 185–191 (2005).

15 Shimada, T. et al. Intracellular disruption of mitochondria in a living HeLa cell with a 76-MHz femtosecond laser oscillator. Opt. Express 13, 9869–9880 (2005).

16 Shi, F. et al. Mitochondrial swelling and restorable fragmentation stimulated by femtosecond laser. Biomed. Opt. Express 6, 4539–4545 (2015).

17 Maxwell, I., Chung, S. & Mazur, E. Nanoprocessing of subcellular targets using femtosecond laser pulses. Med. Laser Appl. 20, 193–200 (2005).

18 Yasukuni, R. et al. Realignment process of actin stress fibers in single living cells studied by focused femtosecond laser irradiation. Appl. Surf. Sci. 253, 6416–6419 (2007).

19 Wang, S. et al. Photoactivation of Extracellular-Signal-Regulated Kinase Signaling in Target Cells by Femtosecond Laser. Laser Photonics Rev. 12, 1700137 (2018).

20 Cheng, P. et al. Direct control of store-operated calcium channels by ultrafast laser. Cell Res. 31, 758–772 (2021).

21 Qi, F., He, H. & Zhu, Y. Neural Development and Repair Induced by Femtosecond Laser Stimulation. ACS Chem. Neurosci. 15, 3106–3112 (2024).

22 Yanik, M. F. et al. Nerve regeneration in Caenorhabditis elegans after femtosecond laser axotomy. IEEE J. Sel. Top. Quantum Electron. 12, 1283–1291 (2006).

23 Yanik, M. F. et al. Functional regeneration after laser axotomy. Nature 432, 822–822 (2004).

24 Dong, B. et al. Spatiotemporally Precise Optical Manipulation of Intracellular Molecular Activities. Adv. Sci. 11, 2307342 (2024).

25 Dong, B. et al. Unleashing precision and freedom of optical manipulation: Software-aided real-time precision opto-control of intracellular molecular activities and cell functions. bioRxiv, 2024.2002. 2009.579709 (2024).

26 Clark, M. G. et al. Real-time precision opto-control of chemical processes in live cells. Nat. Commun. 13, 4343 (2022).

27 Clark, M. G., Gonzalez, G. A. & Zhang, C. Pulse-picking multimodal nonlinear optical microscopy. Anal. Chem. 94, 15405–15414 (2022).

28 Long, G. et al. Targeted tissue ablation with nanosecond pulses. IEEE Trans. Biomed. Eng. 58, 2161–2167 (2011).

29 Oraevsky, A. A. et al. Plasma mediated ablation of biological tissues with nanosecond-to-femtosecond laser pulses: relative role of linear and nonlinear absorption. IEEE J. Sel. Top. Quantum Electron. 2, 801–809 (1996).

30 Venugopalan, V., Guerra III, A., Nahen, K. & Vogel, A. Role of laser-induced plasma formation in pulsed cellular microsurgery and micromanipulation. Phys. Rev. Lett. 88, 078103 (2002).

31 Sims, C. E. et al. Laser− micropipet combination for single-cell analysis. Anal. Chem. 70, 4570–4577 (1998).

32 Watanabe, W. et al. Femtosecond laser disruption of subcellular organelles in a living cell. Opt. Express 12, 4203–4213 (2004).

33 Botvinick, E., Venugopalan, V., Shah, J., Liaw, L. & Berns, M. Controlled ablation of microtubules using a picosecond laser. Biophys. J. 87, 4203–4212 (2004).

34 Sacconi, L., Tolić-Nørrelykke, I. M., Antolini, R. & Pavone, F. S. Combined intracellular three-dimensional imaging and selective nanosurgery by a nonlinear microscope. J. Biomed. Opt. 10, 014002-014002-014005 (2005).

35 Shen, N., Datta, D., Schaffer, C. B. & Mazur, E. Ablation of cytoskeletal filaments and mitochondria in live cells using a femtosecond laser nanoscissor. Mol. Cell. Biomech. 2, 17 (2005).

36 Antunes, F. & Cadenas, E. Estimation of H2O2 gradients across biomembranes. FEBS Lett. 475, 121–126 (2000).

37 Millare, B., O’Rourke, B. & Trayanova, N. Hydrogen peroxide diffusion and scavenging shapes mitochondrial network instability and failure by sensitizing ROS-induced ROS release. Sci. Rep. 10, 15758 (2020).

38 Szechyńska-Hebda, M., Ghalami, R. Z., Kamran, M., Van Breusegem, F. & Karpiński, S. To be or not to be? Are reactive oxygen species, antioxidants, and stress signalling universal determinants of life or death? Cells 11, 4105 (2022).

39 Hockberger, P. E. et al. Activation of flavin-containing oxidases underlies light-induced production of H2O2 in mammalian cells. Proc. Natl. Acad. Sci. U. S. A. 96, 6255–6260 (1999).

40 Mahapatra, S., Ma, S., Dong, B. & Zhang, C. Quantification of cellular phototoxicity of organelle stains by the dynamics of microtubule polymerization. VIEW 5, 20240013 (2024).

