## Supplementary figures for "Real-Time and Site-Specific Perturbation of Dynamic Subcellular Compartments Using Femtosecond Pulses"

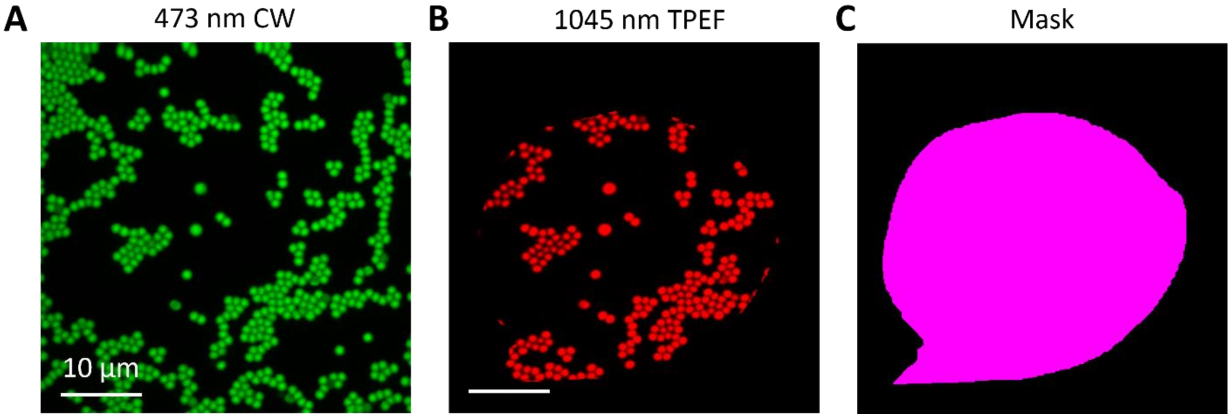


**Figure S1.** Characterization of the fs-RPOC system. (A) Fluorescent microparticles (1 µm in diameter) excited by a 473 nm CW laser. (B) The same field of view as in panel A, with fluorescent microparticles excited by a 1045 nm fs laser via two-photon absorption. (C) RPOC software-generated mask used for TPEF imaging and RPOC testing. The fluorescence signals in panels A and B are detected from the same signal channel.

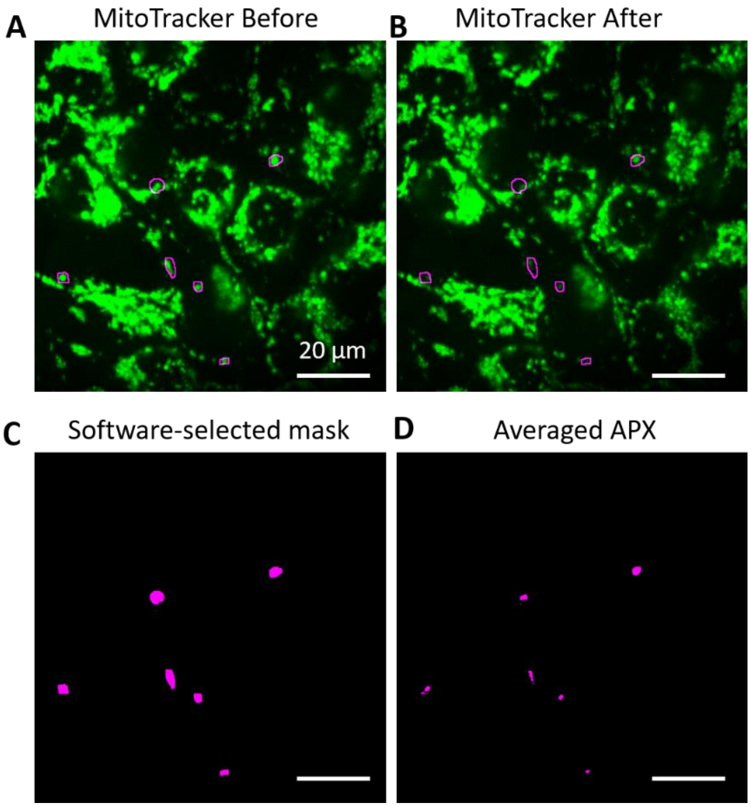


**Figure S2.** fs-RPOC enabled single- and sub-organelle microsurgery. (A, B) MitoTracker signals before and after RPOC. The RPOC-software-selected ROIs are outlined in magenta. (C) RPOC software-selected ROIs. (D) Averaged APX automated selected within the selected ROIs by comparator circuits.

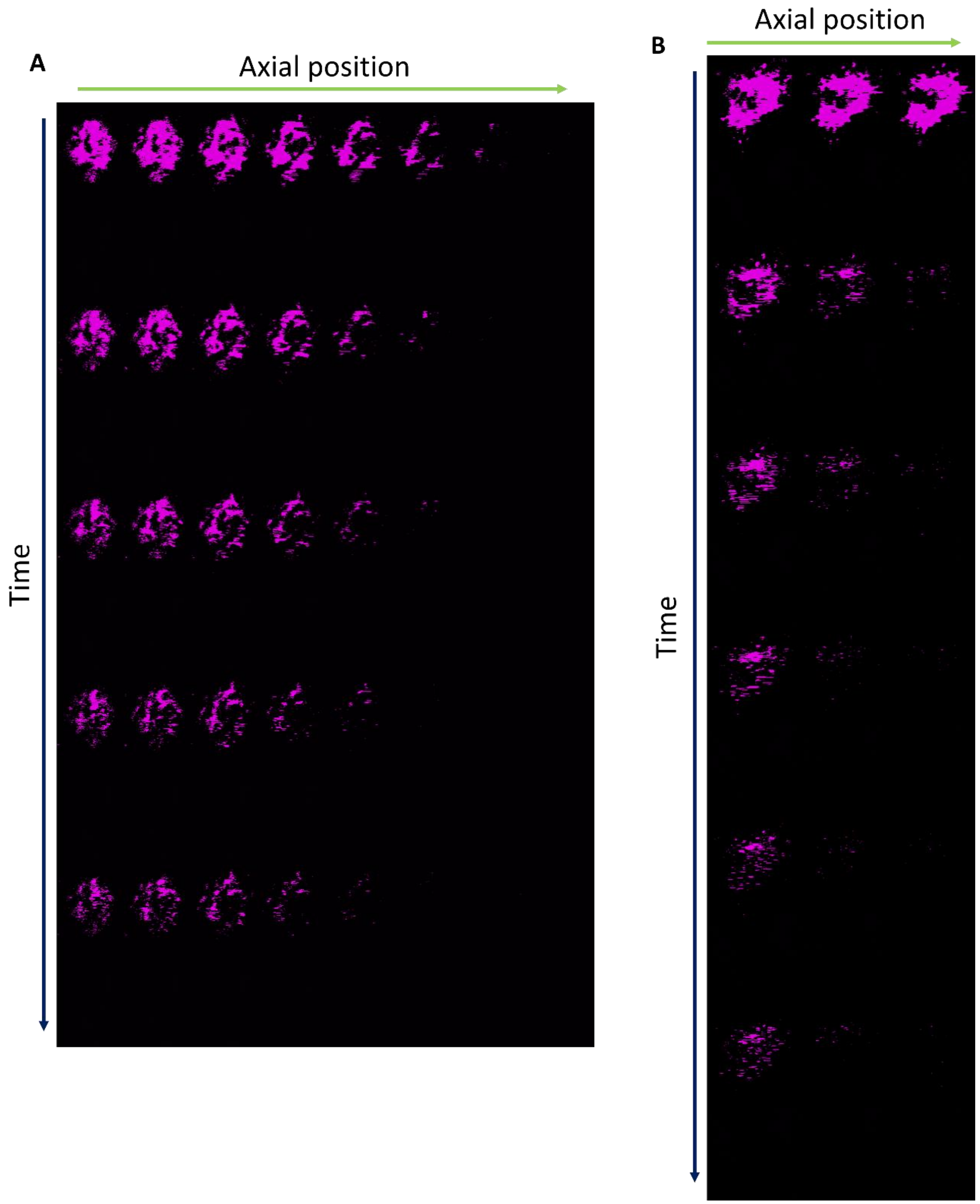


**Figure S3.** APX measurements across different depths and treatment durations corresponding to Figure 3. (A) APXs for the treatment of the 8 µm layer shown in Figures 3F–J. A total of 8 layers in axial direction are scanned at each time point for 5 time points. The APX images are cropped from the original image similar to Figure 3G. (B) APXs for the treatment of the 3 µm layer shown in Figures 3K–O. A total of 3 layers in axial direction are scanned at each time point for 6 time points. The APX images are cropped from the original image similar to Figure 3L. Treatment is performed at 2.6 s per frame.

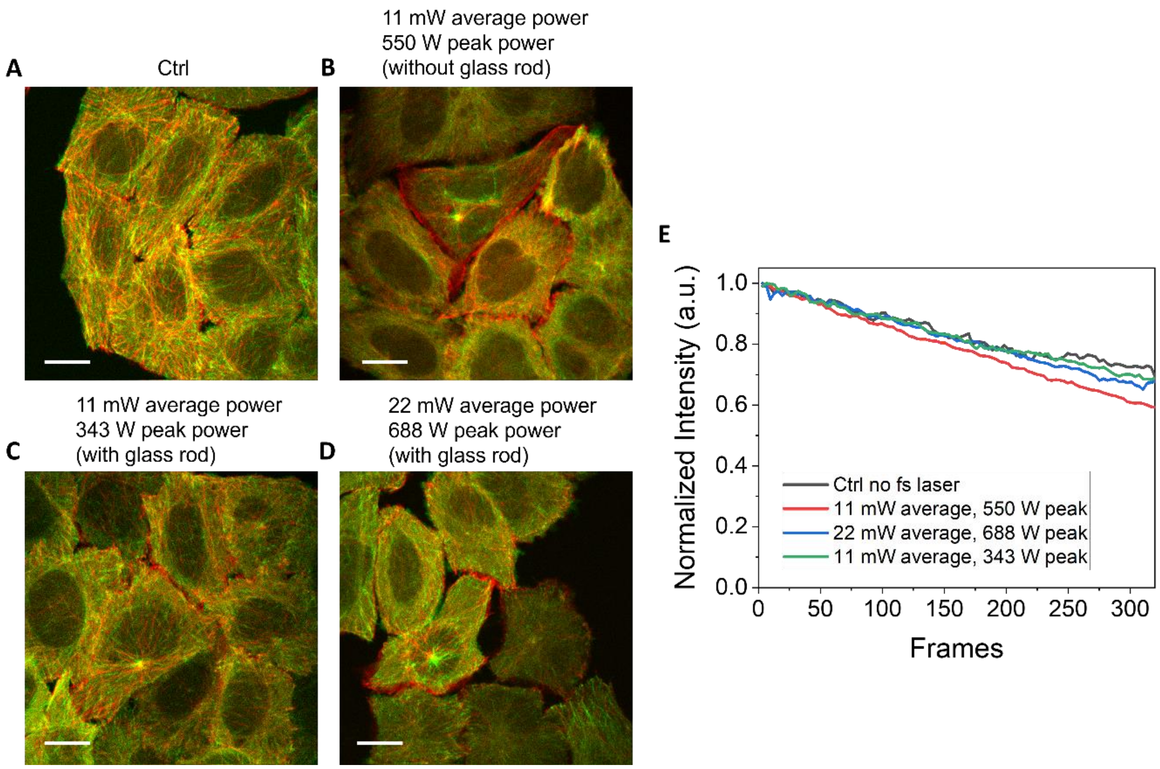


**Figure S4.** Interaction of lower peak power 1045 nm femtosecond laser with cells. (A) EB3-EGFP cells imaged with a 450 μW 473 nm laser for 320 s. The red and green contrasts represent the averaged images from the first and last five frames, respectively. (B–C) Similar to panel A, but with cells exposed to different average and peak powers of a 1045 nm fs laser. (E) Averaged EB3-EGFP signals from cells under different conditions shown in panels A–D.

### Supplementary Videos

Video S1. Real-time APX and MitoTracker signal changes for cells displayed in Figures 3B,C.

Video S2. Time-lapse EB3-EGFP signals for cells in Figure 4B.

Video S3. Time-lapse EB3-EGFP signals for cells in Figure 4C.

Video S4. Time-lapse EB3-EGFP signals for cells in Figure S4.

Video S5. Time-lapse EB3-EGFP signals for cells in Figure 5A.
